# Spontaneous symmetry breaking of cooperation between species

**DOI:** 10.1101/2024.05.27.596113

**Authors:** Christoph Hauert, György Szabó

**Affiliations:** Department of Mathematics, University of British Columbia, 1984 Mathematics Road, Vancouver B.C. Canada, V6T 1Z2; Department of Zoology, University of British Columbia, 6270 University Boulevard, Vancouver B.C. Canada, V6T 1Z4; Institute of Technical Physics and Materials Science, Centre for Energy Research, P.O. Box 49, 1525 Budapest, Hungary; Institute of Evolution, Centre for Ecological Research, Konkoly-Thege M. út 29-33, 1121 Budapest, Hungary

**Author notes:** The authors contributed equally.

**Keywords:** social dilemmas, cross species interactions, phase transitions, structured populations, prisoner’s dilemma

## Abstract

In mutualistic associations two species cooperate by exchanging goods or services with members of another species for their mutual benefit. At the same time competition for reproduction primarily continues with members of their own species. In *intra*-species interactions the prisoner’s dilemma is the leading mathematical metaphor to study the evolution of cooperation. Here we consider *inter* -species interactions in the spatial prisoner’s dilemma, where members of each species reside on one lattice layer. Cooperators provide benefits to neighbouring members of the other species at a cost to themselves. Hence, interactions occur across layers but competition remains within layers. We show that rich and complex dynamics unfold when varying the cost-to-benefit ratio of cooperation, *r*. Four distinct dynamical domains emerge that are separated by critical phase transitions, each characterized by diverging fluctuations in the frequency of cooperation: *(i)* for large *r* cooperation is too costly and defection dominates; *(ii)* for lower *r* cooperators survive at equal frequencies in both species; *(iii)* lowering *r* further results in an intriguing, spontaneous symmetry breaking of cooperation between species with increasing asymmetry for decreasing *r*; *(iv)* finally, for small *r*, bursts of mutual defection appear that increase in size with decreasing *r* and eventually drive the populations into absorbing states. Typically one species is cooperating and the other defecting and hence establish perfect asymmetry. Intriguingly and despite the symmetrical model setup, natural selection can nevertheless favour the spontaneous emergence of asymmetric evolutionary outcomes where, on average, one species exploits the other in a dynamical equilibrium.

## 1 Introduction

The evolution and maintenance of cooperation ranks among the most fundamental questions in biological, social and economical systems (Traulsen and Glynatsi, 2023). Cooperators provide benefits to others at a cost to themselves. Cooperation is a conundrum because on the one hand everyone benefits from mutual cooperation but on the other hand individuals face the temptation to increase their personal gains by defecting and withholding cooperation. This generates a conflict of interest between the group and the individual, which is termed a social dilemma (Axelrod, 1984; Dawes, 1980; Hauert et al., 2006). The prisoner’s dilemma is the most popular mathematical metaphor of a social dilemma and theoretical tool to study cooperation through evolutionary game theory (Maynard Smith, 1982; Nowak, 2006; Tanimoto, 2015). In the simplest and most intuitive instance of the prisoner’s dilemma, the donation game (Sigmund, 2010), two players meet and decide whether to cooperate and provide a benefit *b* to their interaction partner at a cost *c* (*b* > *c* > 0) or to defect, which entails no costs and provides no benefits. If both cooperate each gets *b* − *c* but if one defects then the defector gets *b* and shirks the costs of cooperation, while the cooperator is stuck with the costs −*c*. Finally if both defect, no one gets anything. The interaction is conveniently summarized in a payoff matrix (for the row player):

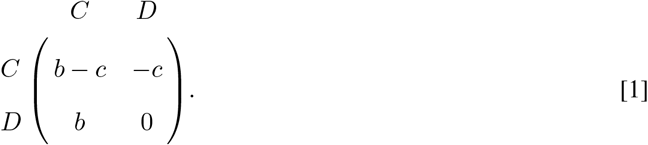

Thus, both players prefer mutual cooperation over mutual defection, yet defection yields the higher payoff regardless of what the partner does and hence the conflict of interest arises. Various mechanisms have been identified that are capable of supporting cooperation in interactions within species. This includes repeated interactions, reputation, punishment or optional participation (Hauert et al., 2002; Nowak and Sigmund, 2005; Sigmund et al., 2001; Trivers, 1971). In structured populations limited local interactions can promote cooperation (Allen et al., 2017; Ohtsuki et al., 2006; Roca et al., 2009; Szabó and Fáth, 2007) and results in complex dynamics and critical phenomena (Szabó and Hauert, 2002).

Another, equally important form of cooperation concerns symbiotic interactions between species that benefit both. Such mutualistic species associations abound in nature with intriguing implications and broad applications. For example, in conservation biology in the context of coral bleaching (the symbiosis of polyps and their algae breaks down, (Wooldridge, 2010)), or pollination (symbiotic interactions of insects and plants, (Bronstein, 2015)), in agriculture (nitrogen fixing microbes living in root nodules of plants, (Kiers et al., 2003)), or pest control (manipulation of microbial symbionts of insects (Douglas, 2007)), as well as in treating disease and supporting health (through interventions in microbiomes associated with animals, including humans (Clemente et al., 2012)).

The problem of cooperation is exacerbated for actions that bestow benefits not just to other individuals but to those of another species (Sachs and Simms, 2006; Werner et al., 2018). In particular it is crucial to clearly distinguish between the acts of cooperation of an individual and mutualistic interactions between individuals. Accomplishing mutually beneficial interactions requires coordination of cooperative acts between species. Moreover, inter-species interactions introduce various additional dimensions that impact evolutionary trajectories. For example, *(i)* interactions (mostly) occur between species, whereas competition (mostly) happens within species, or, *(ii)* generation times between species can differ by orders of magnitude with profound ecological implications, or, *(iii)* body size can hugely differ such that entire populations of one species live on or in a single individual of the other species. Here we focus on the simplest setup with two identical species of equal population size each following the same ecological and evolutionary updating process.

The spontaneous symmetry breaking reported here represents new territory for social interactions in evolutionary biology, while relating to well-known critical phenomena in statistical physics. This is exemplified by the spontaneous magnetization in the Ising model through symmetry breaking in the absence of an external magnetic field. This process is closely related our observations despite having further degrees of freedom. The details of critical phase transitions may be only of limited relevance to biological or social settings in nature. However, the fact that the underlying dynamics has the potential to drive unbounded fluctuations, or give rise to diverging spatial correlations, may have significant implications for the study and management of biological or social systems.

### 1.1 Cooperation between species

Cooperation between species emphasizes *inter-species interactions* as opposed to *intra-species competition*. This is of the essence for all symbiotic species associations (Doebeli and Knowlton, 1998; Frederickson, 2017). In inter-species interactions it is of crucial importance to distinguish between cooperation – costly acts of individuals that benefit others – and mutualistic interactions, where both parties benefit from the interaction. For the inter-species donation game cooperators of one species, *X*, provide benefits to members of another species, *Y*, at a cost to themselves. In well-mixed populations, where members of one species interact with random individuals of the other species, the corresponding two-species replicator equation, for the frequencies of cooperators *x* in one species and *y* in the other, is given by

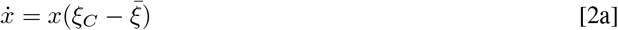

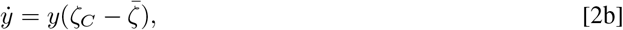

where *ξ*_*C*_ = *yb*_*x*_ − *c*_*x*_ and *ζ*_*C*_ = *xb*_*y*_ − *c*_*y*_ denote the expected payoffs for cooperators in species *X* and *Y* with costs *c*_*x*_, *c*_*y*_ and benefits *b*_*x*_, *b*_*y*_, respectively. Defectors equally reap the benefits but bear no costs, *ξ*_*D*_ = *yb*_*x*_ and *ζ*_*D*_ = *xb*_*y*_, respectively. Note, the benefits to either species depend on the frequency of cooperators in the other species, who also bears the costs of cooperation. The average payoff to each species is 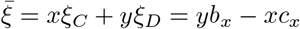 and 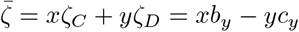, respectively.

Eq. 2 reduces to 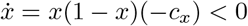 and 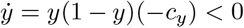 such that in both species cooperators dwindle and eventually go extinct. Interestingly, this happens regardless of the magnitude of foregone benefits, *b*_*x*_, *b*_*y*_. This is a feature of the donation game, or more generally, of additive games (Nowak and Sigmund, 1990). Thus, unsurprisingly the demise of cooperation in inter-species interactions remains unchanged.

For simplicity we assume that species *X* and *Y* face the same donation game, with costs *c* = *c*_*x*_ = *c*_*y*_ and benefits *b* = *b*_*x*_ = *b*_*y*_. Thus, *X* and *Y* are interchangeable labels. Rescaling reduces the interaction in Eq. 1 to a single parameter:

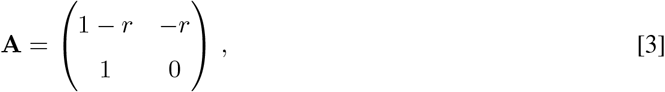

where *r* = *c/b* denotes the cost-to-benefit ratio with 0 <*r*< 1.

In order to delineate effects of spatial dimensions on cooperation in mutualistic interactions, we consider two parallel, square lattice layers with *N* = *L* × *L* sites each, where *L* refers to the linear lattice dimension. Each site is occupied by one individual with *k* = 4 neighbours under periodic boundary conditions. One layer represents species *X* and the other species *Y* (Doebeli and Knowlton, 1998; Ezoe and Ikegawa, 2013): individuals in each layer interact with their *k* + 1 nearest neighbours on the *other* layer but compete with *k* neighbours in their *own* layer, Fig. 1. Intriguing examples of multiple species interacting in spatial settings naturally arise in microbial communities and biofilms, in particular (Kreft, 2004; Schuster et al., 2010) and may even give rise to actual physical layering of mutualistic partner species (Momeni et al., 2013).

**Fig. 1.**
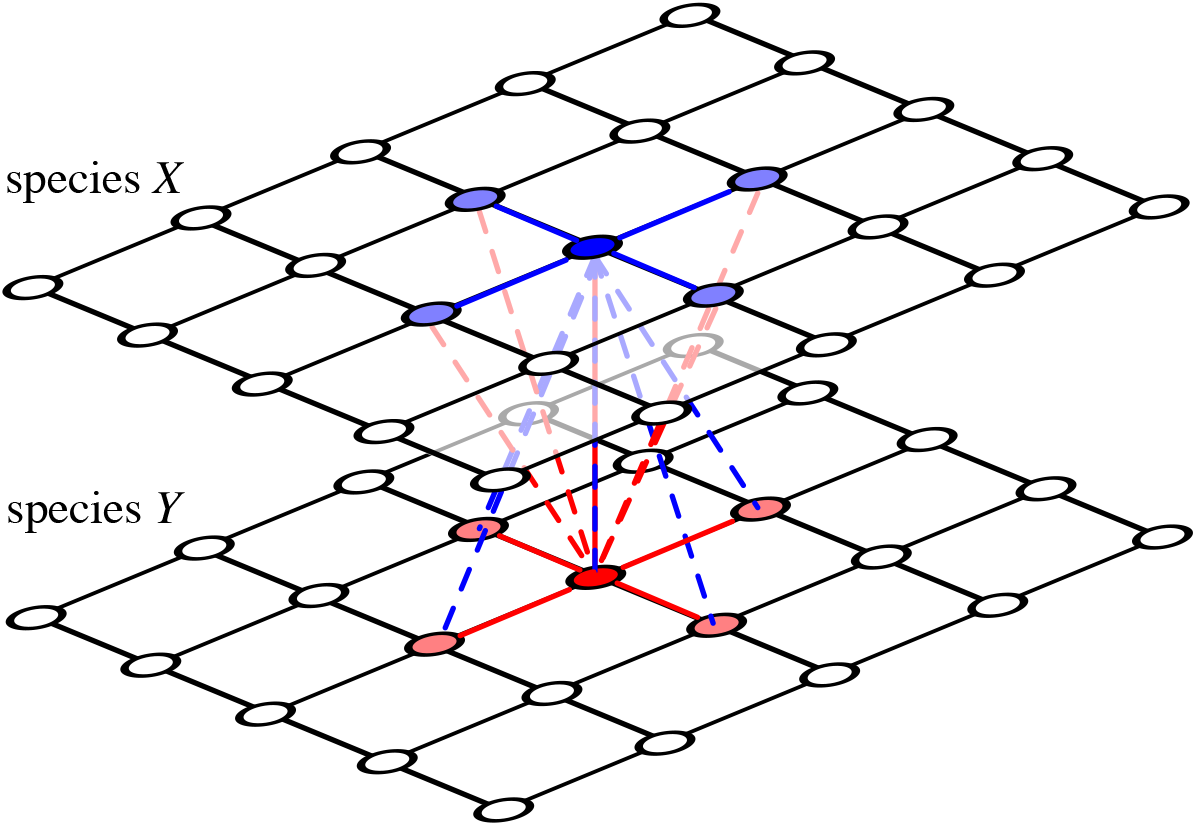
(colour online) On lattices with *k* = 4 neighbours, an individual in species 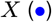 interacts (dashed lines) with its *k* + 1 neighbours in the other species *Y* (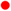 and 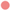) but competes (solid blue lines) with its *k* neighbours in its own species 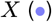. Analogously, a member of species 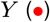 interacts with their neighbours in species *X* (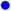 and 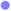) and competes with neighbours of their own species 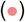. Click image for interactive online simulations of the two-layer donation game with *r* = 0.02.

For an individual *i* in layer *a* ∈ {−1, +1} cooperation, *C*, and defection, *D*, are represented by unit vectors

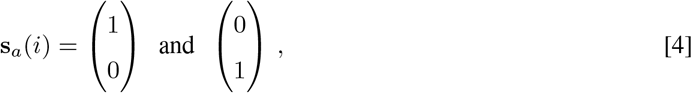

respectively. The (accumulated) payoff to individual *i* is given by

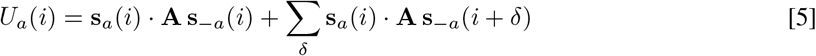

where the summation runs over the *k* neighbours, *i* + *δ*, on the *other* layer, −*a*. In contrast, competition arises among the *k* neighbours on the *same* layer. A randomly chosen focal individual *i* competes with one of its neighbours *j*. With probability

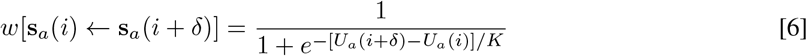

the focal individual adopts (imitates) the neighbours’ strategy or, equivalently, the neighbour produces a clonal offspring, which successfully displaces the focal individual. *K* determines the noise in the updating process. Reproduction (imitation) invariably generates positive assortment of strategic types among neighbours within layers but the payoffs from interactions can drive positive or negative assortment of strategic types between layers. For *K* ≫ 1, imitation (reproduction) reduces to a coin toss (*w* → 1*/*2 for *K* → ∞). In this limit the configurations in each layer are uncorrelated and payoffs irrelevant. In contrast, with little noise, *K* ≪ 1, selection is strong and even marginally better performing neighbours are almost certainly imitated (reproduce successfully). Interactions can now indirectly generate strong correlations of strategic types between layers. This updating admits four absorbing states where each of the two layers are homogeneous *C* or *D* states.

Eq. 6 creates strong correlations among neighbours within layers through imitation or reproduction and weaker correlations across layers, which are mediated by the performance of each strategy in interactions. For large *r* correlations within and across layers are too weak and cooperation vanishes just as in the well-mixed scenario or in *intra*-species spatial games. However, for sufficiently small *r*, the spatial structure enables cooperators to form clusters and correlations between lattices are capable of supporting cooperation. This is similar to intra-species donation games (Allen et al., 2017; Ohtsuki et al., 2006; Szabó and Fáth, 2007), but with much richer and indeed surprising dynamical behaviour.

## 2 Dynamical scenarios

The average frequency of cooperators, 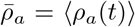, reveals a sequence of distinct dynamical regimes as a function of *r. ρ*_*a*_(*t*) denotes the cooperator frequency at time *t* in layer *a* and the averaging runs over a sampling time of *τ*_*s*_ Monte Carlo (MC) steps (*N* individual updates each) after a relaxation time *τ*_*r*_, see Fig. 2. Note, large lattice sizes are required to prevent accidental absorption due to large (indeed at times diverging), systematic dynamical fluctuations. The relatively short relaxation times of *τ*_*r*_ = 10^5^ for the baseline simulations with *L* = 800 in Figs. 2, 3 are justified by using the equilibrated state of simulations with adjacent *r* values as the initial configuration. As a consequence, only small changes are required to attain the new equilibrium state. In the vicinity of the critical transitions this is insufficient and more sophisticated methods that aggregate data from many simulation runs are needed to determine the critical points as well as the equilibrium behaviour in their vicinity.

**Fig. 2.**
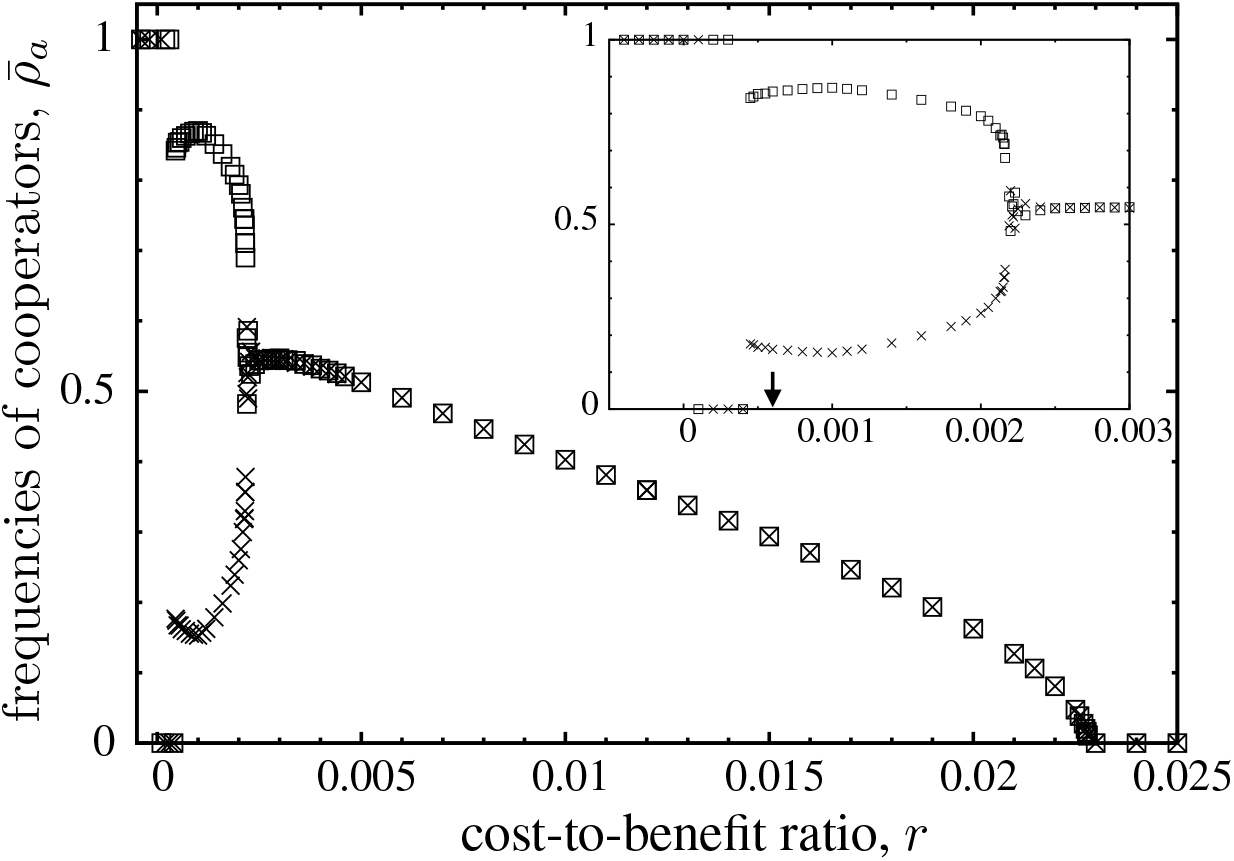
Frequency of cooperation 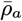 in each layer (□ and ×, respectively) as a function of the cost-to-benefit ratio *r* for *K* = 0.1. The inset enlarges the region of spontaneous symmetry breaking and the arrow indicates the *r* for which a burst of defection is illustrated in Fig. 11. Simulations for lattices with linear size *L* = 800, after a relaxation time *τ*_*r*_ = 10^5^ and averages over *τ*_*s*_ = 10^5^ MC steps and up to *L* = 1800, *τ*_*r*_ = 2 · 10^6^ and *τ*_*s*_ = 10^7^ in the vicinity of critical points.

**Fig. 3.**
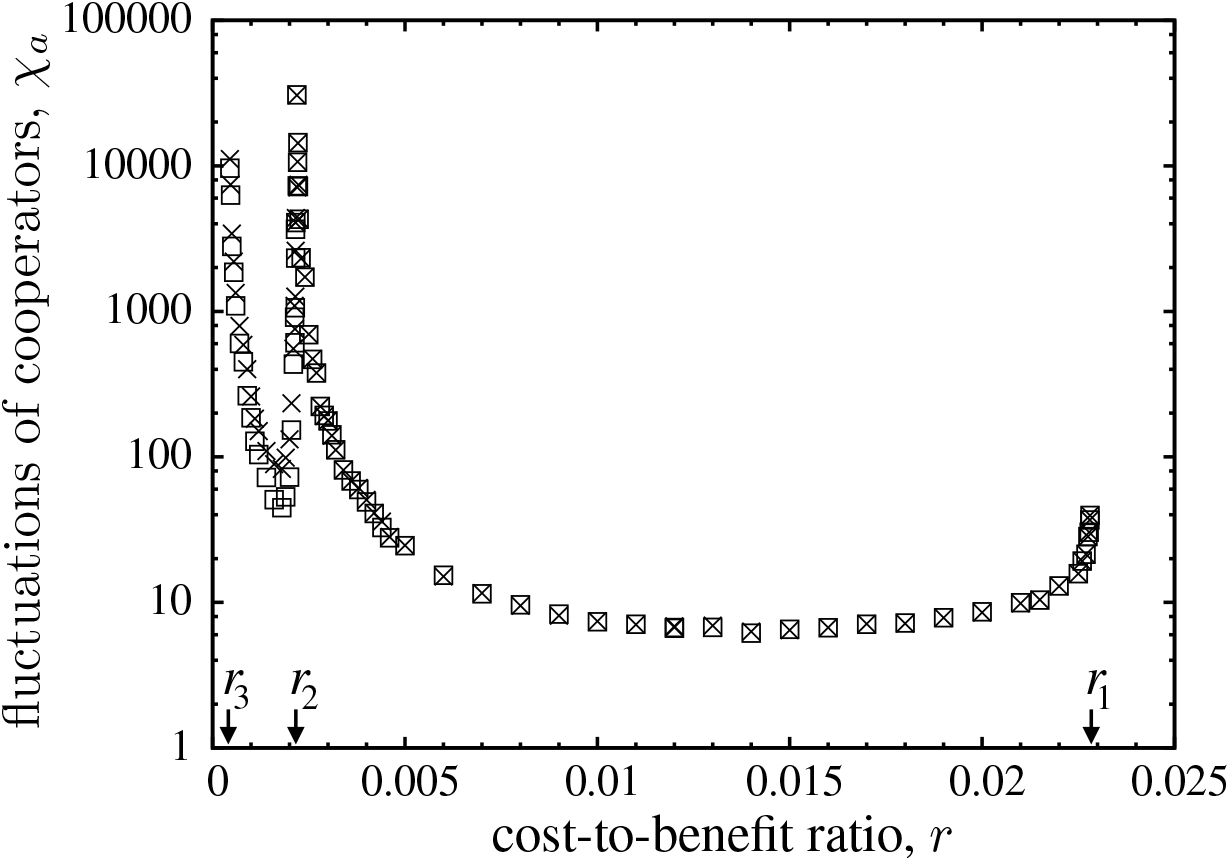
Fluctuations of the frequencies of cooperation, *χ*_*a*_, in each layer (□ and ×) for *K* = 0.1. This reveals three critical phase transitions: *r*_1_, *r*_2_ and *r*_3_, which are characterized by their asymptotic power law scaling of diverging fluctuations. Simulation parameters as in Fig. 2.

First, for *r* < 0 the social dilemma disappears and the interaction, Eq. 3, turns into a harmony game in which *C* is dominant (mutual cooperation is the sole Nash equilibrium). More precisely, with *k*_*c*_ cooperating neighbours the payoff to a defector is *k*_*c*_ and always lower than that to a cooperator, *k*_*c*_ − *k r*. Hence, regardless of the strategies in the other layer, *C* players are better off, such that both lattices evolve to homogeneous cooperation. Even though this seems a natural outcome, it is worth noting that this actually contrasts with results from intra-species interactions. On a single lattice solitary, or small groups of defectors can survive fo −1*/k* < *r*, where *k* is the number of interaction partners. An isolated defector achieves the maximum score of *k*, while each one of its cooperating neighbours gets *k*(1 − *r*) − 1. Thus for −1*/k* < *r* < 0 isolated defectors always outperform cooperators. As *r* increases pairs or small clusters of defectors may prevail too. This is surprising because it renders defection an act of spite (Hamilton, 1970), where defectors *lower* the benefit of others at a cost to themselves. Spiteful behaviour only works in small groups or, in spatial settings, on a local scale (Smead and Forber, 2013; Szabó and Fáth, 2007). Fortunately such behaviour is unable to spread because the losses incurred from mutual defection cannot compensate for the additional benefits of mutual cooperation.

Second, for *r* = 0 the payoff to an individual is independent of its own strategy and hence exploitation (as well as spiteful behaviour) is impossible. Since *CC* interactions benefit individuals in both layers and because exploitation is not an issue, the frequency of cooperators keeps increasing. Once defectors become extinct in one layer, both types in the other layer receive the same payoff. Their dynamics recovers the two-dimensional voter model (Liggett, 1991), which exhibits very slow (logarithmic) domain coarsening (Ben-Naim et al., 1996) towards a homogeneous absorbing state with either *C* or *D* players.

Finally, for *r* > 0 the social dilemma in Eq. 3 is in effect and *D* dominates (mutual defection is the sole Nash equilibrium). However, in spatial settings homogeneous defection results only for sufficiently large *r* > *r*_1_ ≈ 0.02283(3). For smaller *r*, the limited local interactions suffice to protect cooperators. The formation of clusters reduces exploitation and increases interactions with other cooperators to make up for losses against defectors. Spatial assortment enables co-existence. More specifically, four dynamical regimes exist that are separated by critical phase transitions. In order to determine the boundaries of the dynamical domains at *r*_1_, *r*_2_, and *r*_3_, in descending order of *r*, as well as to classify the accompanying phase transitions, we consider the fluctuations, *χ*_*a*_, in layer *a*:

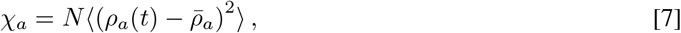

where *χ*_*a*_ becomes constant in the limit *N* → ∞ (Stanley, 1999) and *χ*_*a*_ = 0 holds for homogeneous states. The scaling of *χ*_*a*_ as *r* approaches a critical phase transition helps to identify the corresponding universality class. The large fluctuations that accompany and indeed characterize the critical phase transitions, require significantly larger lattices (up to *L* = 1800 with *τ*_*s*_ > 10^7^, *τ*_*r*_ = 2 · 10^6^), see Fig. 3. The four dynamical domains are:

i. *r* > *r*_1_: defection dominates. For high costs or low benefits, spatial correlations are insufficient to support cooperation and both populations evolve towards the absorbing *DD* state. The outcome is the same as in well-mixed populations, see Eq. 2.
ii. *r*_2_ < *r* < *r*_1_: cooperators and defectors co-exist. The frequency of cooperators in the dynamical equilibrium is equal in both layers and increases for decreasing *r*.
iii. *r*_3_ < *r* < *r*_2_: spontaneous symmetry breaking. Cooperators and defectors continue to co-exist in a dynamical equilibrium but with essentially complementary strategy frequencies in the two layers.
iv. 0 < *r* < *r*_3_: relaxation into asymmetric absorbing states *CD* or *DC*. The emergence, growth and elimination of homogeneous domains of defection (*DD* regions) drive spikes (or bursts) in defector frequencies.

The evolution towards absorbing states for very small *r* is a long-time process preceded by a spatial phase separation. For increasing *r* the size of these bursts decreases and are no longer able to maintain global synchronization across and between layers. Instead, the bursts break the symmetry between the two layers resulting in different average frequencies of cooperation, 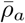, in each layer *a*, see Fig. 2. The asymmetry between layers, 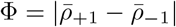, decreases with increasing *r* until cooperation in both layers is the same, 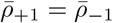 Further increases of *r* monotonously decrease 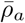 until eventually cooperators go extinct for *r* > *r*_1_.

### 2.1 Directed Percolation

Near the extinction threshold, *r*_1_, isolated clusters of cooperators perform a branching and annihilating random walk with almost perfect correlations between lattices. The characteristic spatial patterns are captured by snapshots of the configuration in each lattice as well as the pair distributions, see Fig. 4. Coordination between lattices is almost perfect with very few *CD* and *DC* pairs. This is of the essence for the survival of cooperation. *CC* interactions are the bulwark against, or at least help to compensate for exploitation by defectors. Interestingly, the emerging spatial configurations are very similar to those in the intra-species donation game on a single lattice (Szabó and Tőke, 1998).

**Fig. 4.**
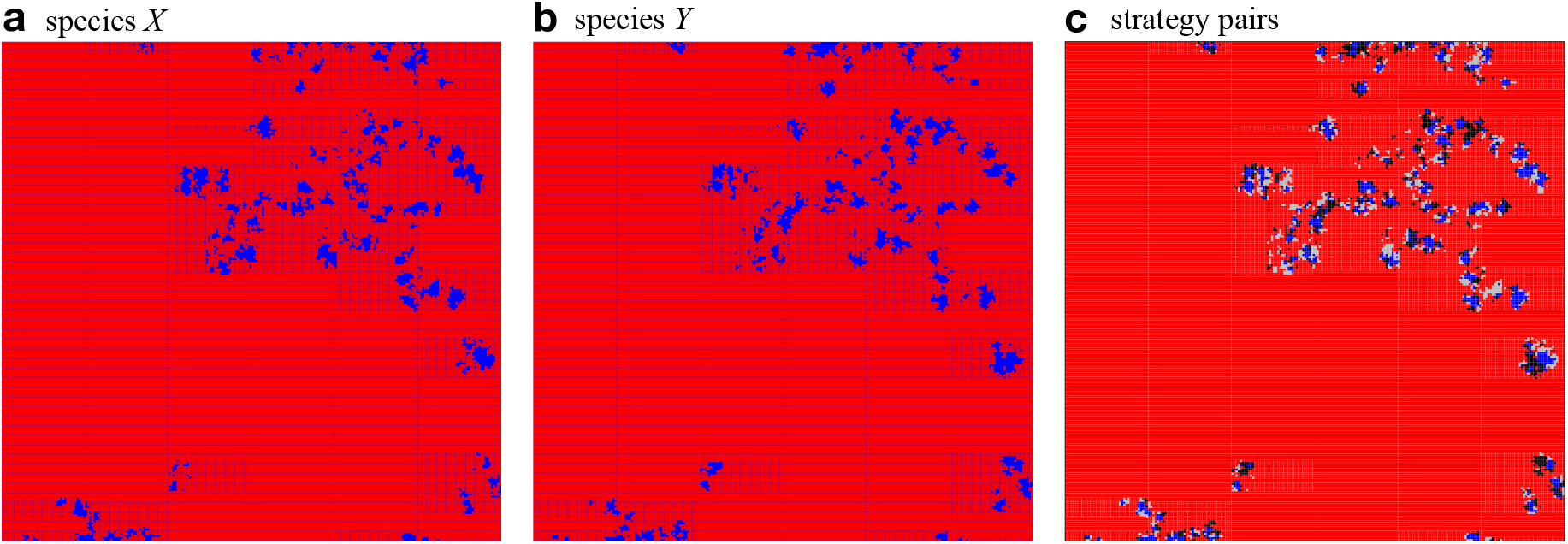
(colour online) The spatial configuration of cooperators, 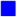, and defectors, 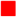 for each species separately, **a** and **b**, as well as for strategy pairs, **c**, with *CC* 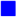, *DD* 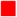, *CD* 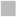 and *DC* ◼ in the symmetric phase close to but below the directed percolation threshold, *r* = 0.0225 < *r*_1_: Strongly correlated clusters of cooperators on both layers perform a coordinated branching and annihilating random walk. Parameters: 200 × 200 region of a 1000 × 1000 lattice with *K* = 0.1. Click image for interactive online simulations.

In agreement with theoretical expectations (Marro and Dickman, 1999; Szabó and Fáth, 2007; Szabó and Hauert, 2002) the critical transition at *r*_1_ clearly belongs to the two-dimensional directed percolation universality class: in both layers the frequency of cooperation scales with 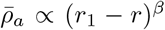 for *β* = 0.580(3) and their fluctuations scale with *χ*_*a*_ ∝ (*r*_1_ − *r*)^−*γ*^ for *γ* = 0.35(1) when *r* → *r*_1_ from below, see Fig. 5.

**Fig. 5.**
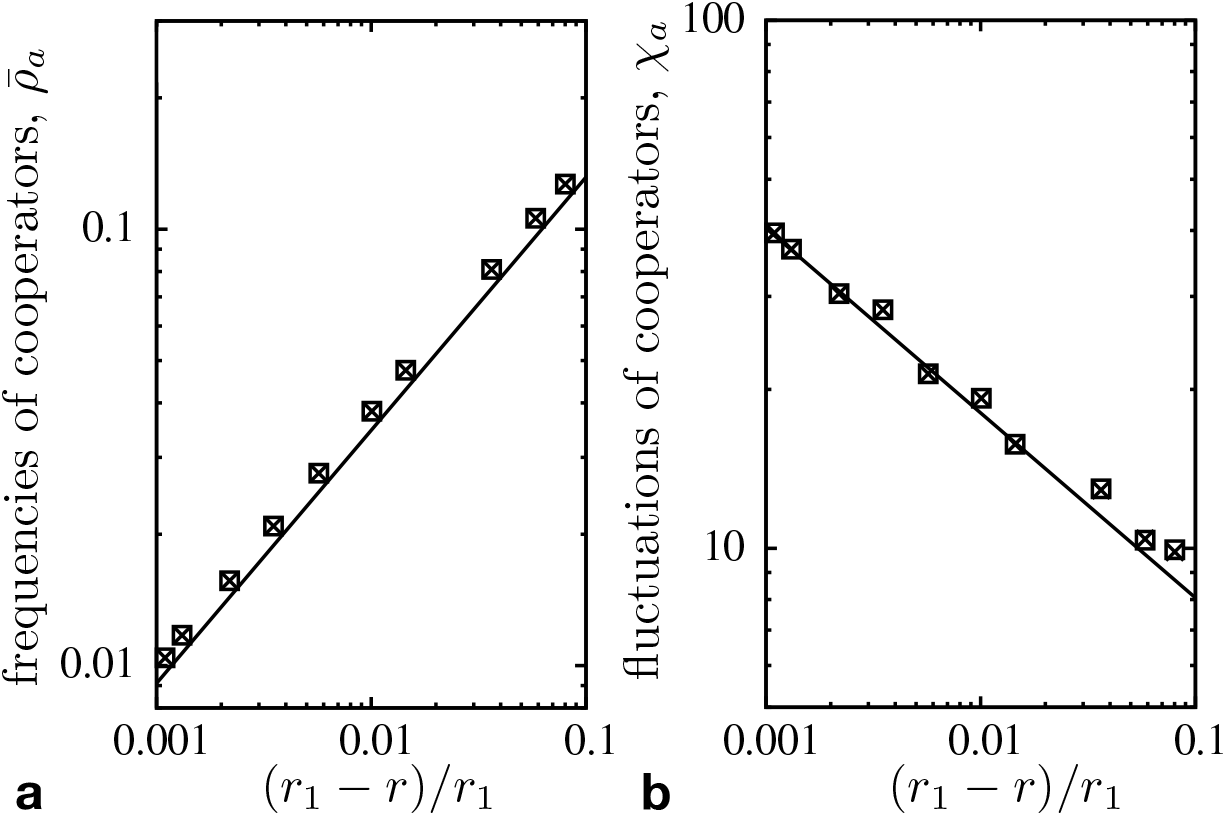
The scaling of **a** 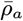 and **b** *χ*_*a*_ versus (*r*_1 ™_ *r*)*/r*_1_ follows the directed percolation universality class (solid lines indicate theoretical expectations).

Decreasing the cost-to-benefit ratio to *r*_2_ < *r* < *r*_1_ with *r*_2_ ≈ 0.002166(3), cooperators persist at equal frequencies in each layer, see Fig. 2 and illustrative snapshots in Fig. 6. In this symmetric phase, the frequency of cooperation gradually increases with decreasing *r*, while the coordination between layers decreases. For *r* close to *r*_2_ the frequency of all four strategy pairs are essentially the same, which supports that configurations are largely uncorrelated.

**Fig. 6.**
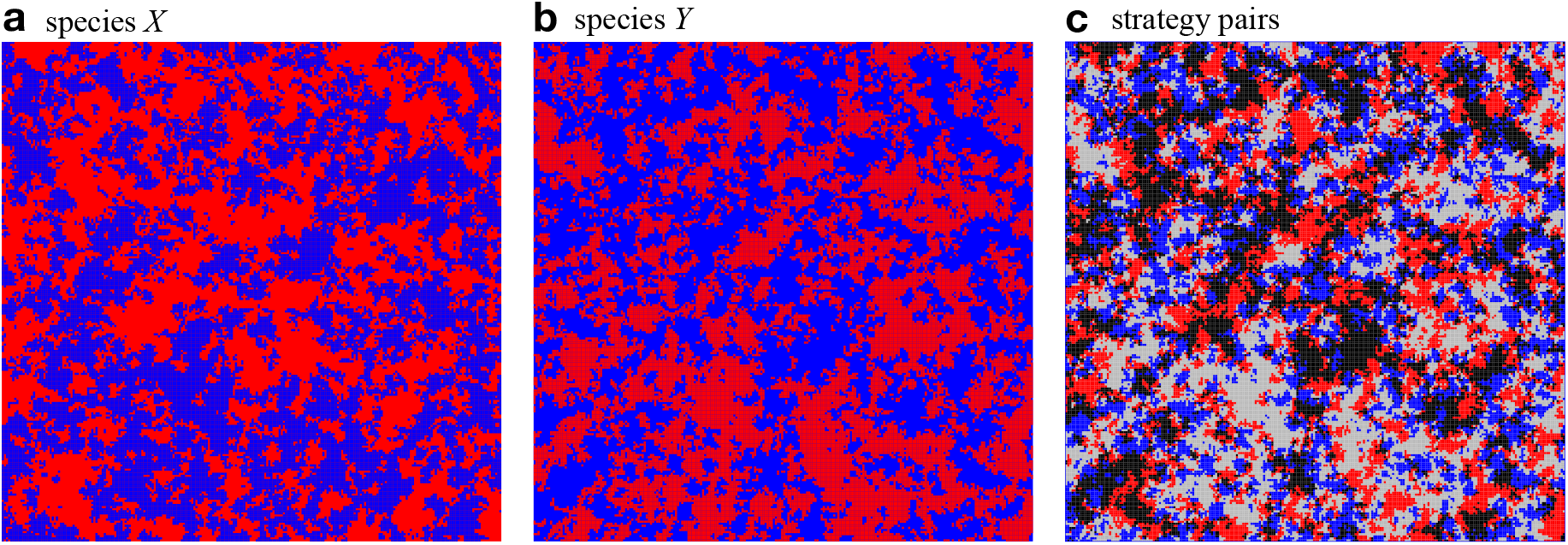
(colour online) The spatial configuration of cooperators, 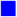, and defectors, 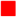 for each species separately, **a** and **b**, as well as for strategy pairs, **c**, with *CC* 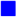, *DD* 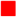, *CD* 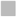 and *DC* ◼ in the symmetric phase close to but above the symmetry breaking, *r* = 0.005 > *r*_2_: Almost identical frequencies of cooperators and defectors establish in both layers. The similar frequencies of all four strategy pairs indicate that the layers are essentially uncorrelated. Switching the labels of species *X* and *Y* in **a, b** merely switches the role of *CD* and *DC* pairs, 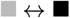. Parameters: 200 *×* 200 region of a 1000 *×* 1000 lattice with *K* = 0.1. Click image for interactive online simulations.

### 2.2 Spontaneous symmetry breaking

Intriguingly, lowering *r* further to *r*_3_ < *r* < *r*_2_ with *r*_3_ ≈ 0.00041(3) results in spontaneous symmetry breaking with different frequencies in each layer, see Fig. 7. The distribution of cooperators and defectors in each layer is almost complementary. Clusters, or regions of cooperators, in one layer are matched by defectors in the other. As a consequence, the lattices consist mostly of *CD* and *DC* pairs. Because of the distinctly different frequencies of cooperation in each lattice, either *CD* or *DC* pairs dominate. However, note that which pair dominates is of no consequence, because species interactions are symmetric, and hence it is merely a consequence of which species is labelled *X* and which *Y*. Defectors in one species manage to exploit and take advantage of cooperators in the other species. However, exploitation is ephemeral and the scale is limited both spatially as well as temporarily.

**Fig. 7.**
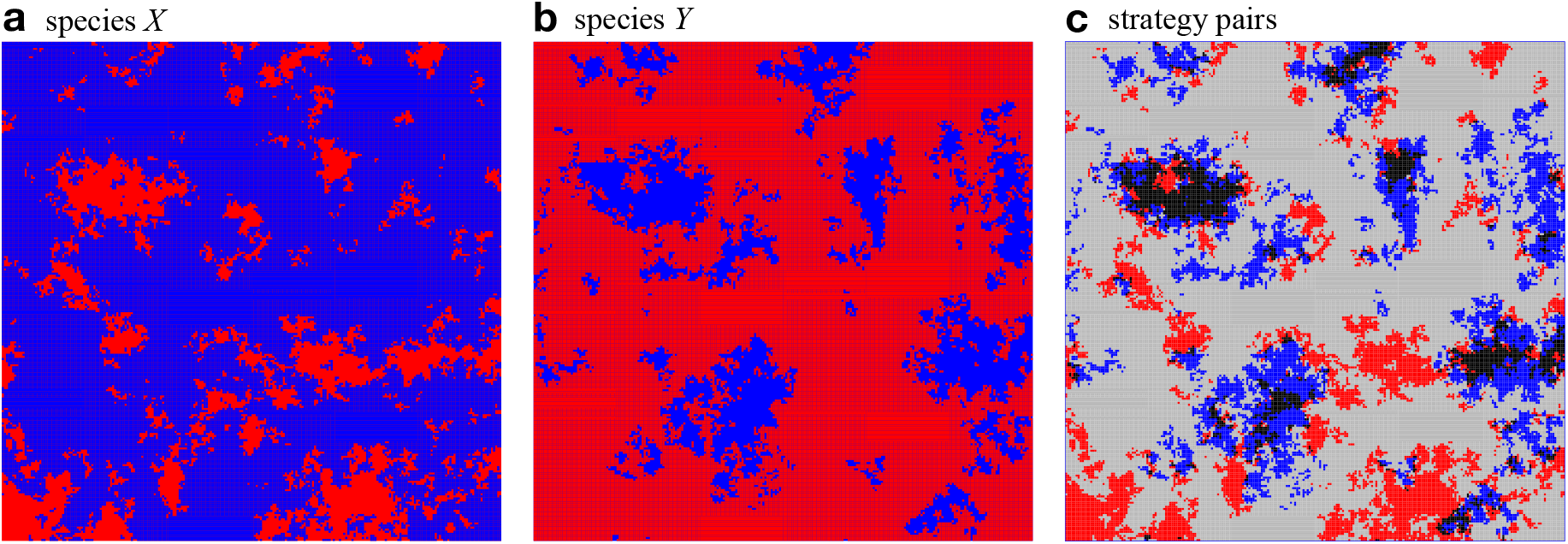
(colour online) Typical spatial strategy distribution in the asymmetric phase with essentially complementary frequencies and distributions of cooperators and defectors in the two layers. Parameters are the same as in Fig. 4 but with *r* = 0.0015. Click image for interactive online simulations.

In the vicinity of the symmetry breaking transition, *r*_2_, technical difficulties can arise in Monte Carlo simulations through the formation of strip-like domain structures that persist for long times at low noise (Lipowski, 1999, 2022; Spirin et al., 2001). Moreover, if both layers are initialized with the same frequency of *C* players, the probability to end up with more *C* players is equal for both layers. Albeit only after a slow domain growing process studied well for Ising models (Furukawa, 1985). Fortunately, such difficulties can be easily prevented by starting from initial configurations with different *C* frequencies in each layer.

Our analysis of the spontaneous symmetry breaking borrows concepts from the Ising model and critical phase transitions (Stanley, 1999). The two-dimensional Ising model exhibits universal features through the power law scaling of the spontaneous magnetization in the absence of an external magnetic field, as well as the accompanying fluctuations when the temperature approaches the critical point. Here we use *r* instead of the temperature as the control parameter and, instead of the magnetization as the order parameter, we use the asymmetry in cooperation between the two layers 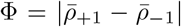. The scaling of Φ and its fluctuations for *r* → *r*_2_, is shown in Fig. 8. The order parameter clearly shows the expected power law behaviour and its fluctuations are in line with the theoretical predictions of |*r* − *r*_2_|^−*γ*^ and *γ* = 7*/*4 within the accuracy of our MC data.

**Fig. 8.**
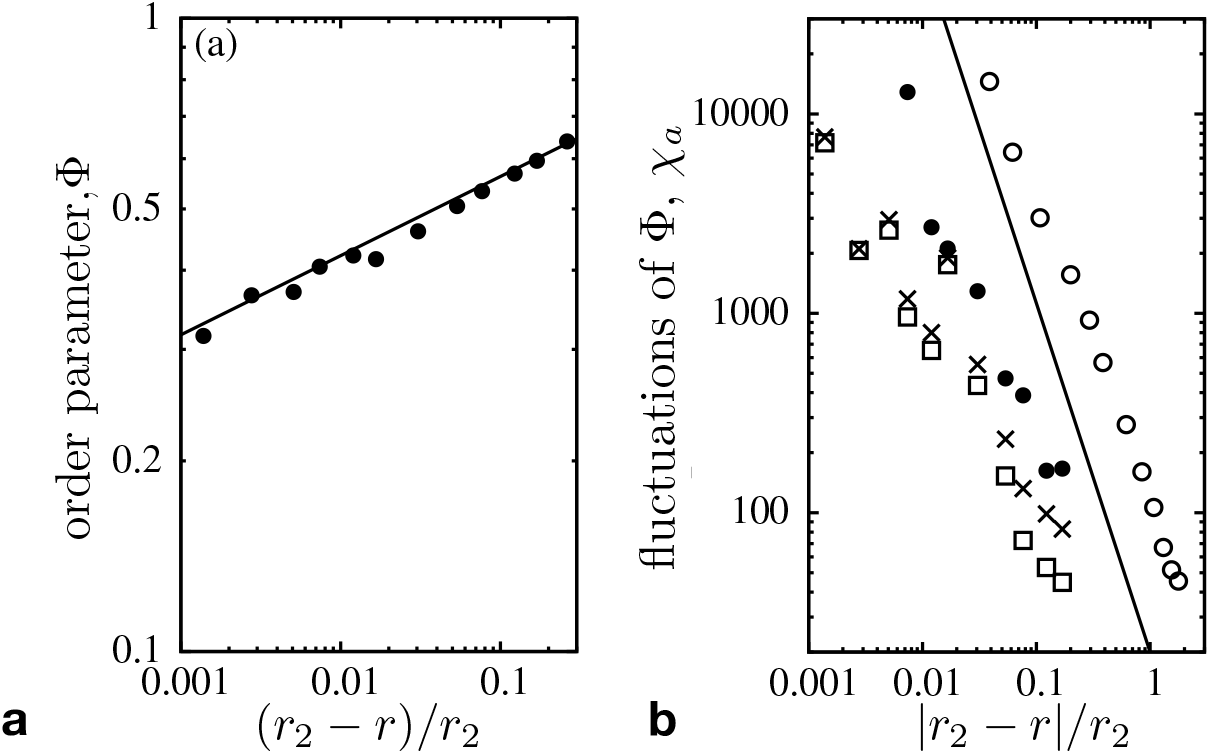
**a** The scaling of the order parameter 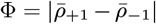 as a function of (*r*_2_ − *r*)*/r*_2_ is in line with the universality class of spontaneous magnetization in the Ising model. **b** The scaling of the fluctuations of Φ (•and ∘) as well as *χ*_*a*_ (□ and *×*) above and below the critical threshold *r*_2_. The Φ data reproduces the theoretical expectations (solid lines with slopes 1*/*8 in **a** and −7*/*4 in **b**) within statistical errors (omitted for clarity).

However, the equivalence to the universal features of Ising type transitions is not obvious. Additional degrees of freedom result from the larger number of possible strategy pairs (*CC, CD, DC*, and *DD*) across the two layers. In order to illustrate the consequences, the scaling of fluctuations of cooperation are shown for each layer separately, see Fig. 8. Disregarding the large uncertainties, Fig. 8b suggest a power law behaviour when approaching the critical point *r*_2_ from either side, albeit with different exponent.

A possible explanation may be related to the anomalous behaviour of the correlation of cooperation between the two layers, which is given by the covariance, *κ*,

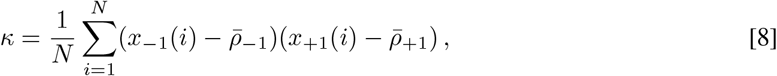

with *x*_*a*_(*i*) = 1, (*x*_*a*_(*i*) = 0) if site *i* in layer *a* is occupied by a cooperator (defector). *κ* exhibits a local maximum for small *r* and a sharp local minimum at *r*_2_, see Fig. 9. Note that the switch in the sign of *κ* does not coincide with the symmetry breaking but happens already at slightly larger *r*.

**Fig. 9.**
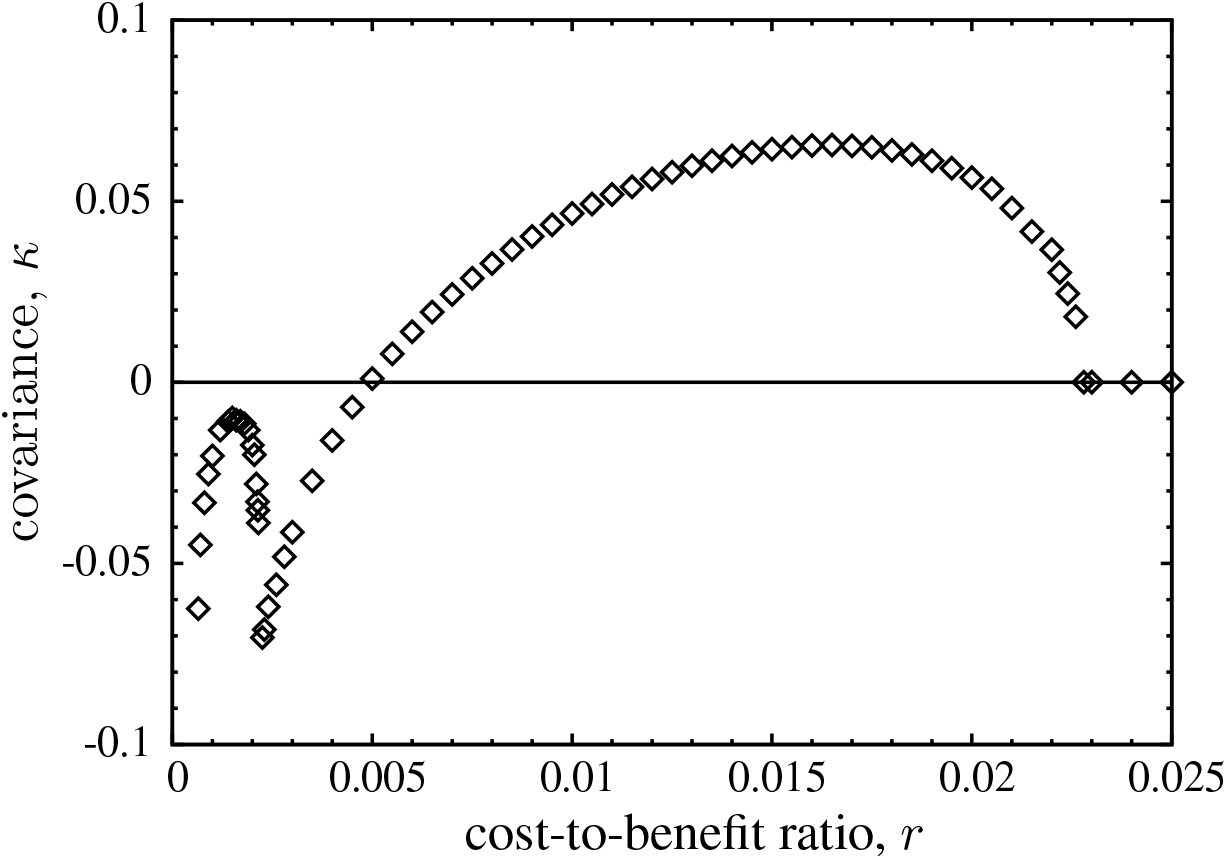
Correlation of cooperators between lattices given by the covariance between layers, *κ*, as a function of *r*. In the symmetric phase correlations are (mostly) positive, while negative in the asymmetric phase. Note the anomalous sharp minimum of *κ* at *r*_2_ and the local maximum at small *r*, which is driven by spikes in mutual defection. Statistical errors become noticeable only for small *r* < 0.00065. This is caused by the possibility of exceedingly long relaxation times that result from the occasional formation of strip-like domain structures.

### 2.3 Bursts of defection

Finally, for 0 < *r* < *r*_3_, the population relaxes into one of its four absorbing states depending on the initial configuration. In fact, the most striking dynamical phenomenon of this model manifests itself in the diverging fluctuations of *C* frequencies, *χ*_*a*_, for *r* → *r*_3_, see Fig. 10. In particular, no theoretical arguments exist to date that explain and justify the observed power law behaviour of *χ*_*a*_ ∝ (*r* − *r*_3_)^−*γ*^ for *r* → *r*_3_ from above with *γ* = 1.33(5). The divergence of *χ*_*a*_ is related to the emergence, growth and elimination of homogeneous *DD* domains. Spikes in *DD* frequencies drive the dynamics of bursts.

**Fig. 10.**
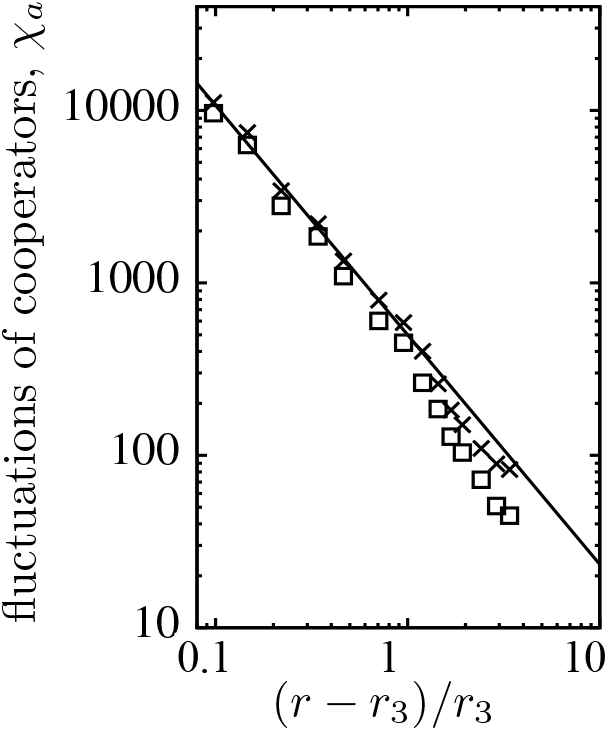
Scaling of *χ*_*a*_ in each layer (□ and *×*) versus (*r* − *r*_3_)*/r*_3_, which is driven by bursts of defection. For reference the solid line shows a power law with exponent −1.33.

In order to illustrate the characteristics of such burst events, we performed MC simulations on a large lattice but focused on the dynamics within a small region, see Fig. 11a. Within this region the *DD* spike is illustrated through a sequence of snapshots, see Fig. 11b-e. First, small islands of *CC* and *DD* pairs move, collide, and disappear at random in a large domain of *CD* pairs (Fig. 11b). These islands are continuously created along the boundary separating large *CD* and *DC* domains (Fig. 11b lower left corner). Most islands eventually go extinct. Only sufficiently large *DD* areas occasionally expand (Fig. 11c). The *DD* domain grows until it collides with a small *CC* island, which invades its territory (Fig. 11d). However, the *CC* domains are not stable against invasion of *CD* or *DC* phases (Fig. 11e). Subsequently, the strategy distribution evolves slowly toward the initial distribution (Fig. 11b).

**Fig. 11.**
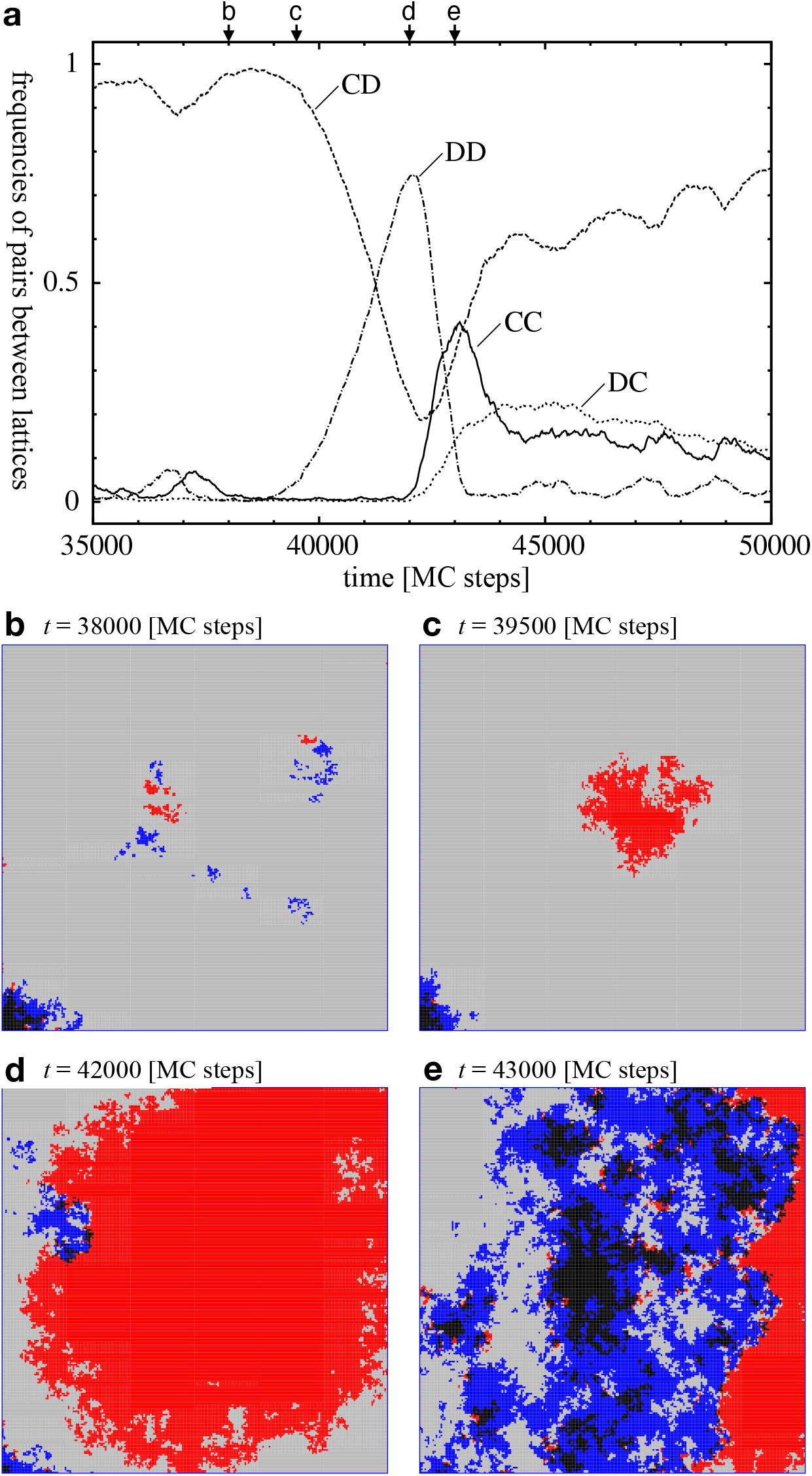
**a** Frequencies of strategy pairs across lattices as a function of time. This illustrates a burst of defection for *r* = 0.0006 within a 240 × 240 region of a 1000 ×1000 lattice. The four arrows mark the times of snapshots. **b**-**e** Snapshots illustrating the spatial distribution of the four strategy pairs between lattices, *CC* 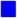, *DD* 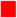, *CD* 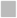 and *DC* ◼, during a *DD* burst within a 240 *×* 240 region of a 1000 *×* 1000 lattice. Click image for interactive online simulations.

This type of bursts are driven by sufficiently large spatial domains of an unstable phase (Hódsági and Szabó, 2019). Evidently, their extension and frequency depends on *r* and other parameters. In particular, bursts become larger and rarer when decreasing *r*, see fig. S1f. Once a burst reaches sizes comparable to the entire lattice, it can drive the evolution towards absorbing states. Interestingly, the absorbing *DD* state is rarely observed because it is prone to invasion by any remaining *CC* islands, which then pave the way for *CD* or *DC* phases to grow and take over. This is in stark contrast to intra-species models where cooperation continuously increases for decreasing *r*, that is as the social dilemma becomes more benign.

Another peculiarity of the burst dynamics is the feedback between their creation and extinction and the spatial configuration of the asymmetric phase: for *r* close to *r*_3_, the asymmetry of cooperation, 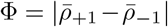 decreases with *r* while the burst sizes increase, see inset in Fig. 2.

For *r* < *r*_3_ the population tends to get absorbed in the fully asymmetric *CD* or *DC* states. Once one layer is homogeneous the dynamics in the other layer is dominated by neutral drift because the difference in payoffs between cooperators and defectors merely amounts to 4*r*. Hence, for small *r*, Eq. 6 essentially reduces to a coin toss with a slight bias in favour of defectors. The probability to fixate in one or the other absorbing state is thus essentially given by the frequency of strategies in the heterogeneous layer at the time the other layer turned homogeneous. For example, if one layer becomes homogeneous in *C*, then the frequency of *C*’s in the other layer is likely < 1*/*2, due to the asymmetry. Thus, the *CD* absorbing state is more likely than *CC*. Conversely, if one layer becomes homogeneous in *D*, then the frequency of *C*’s in the other layer is likely > 1*/*2 and hence the *DC* absorbing state is more likely than *DD*. Finally, it is more likely that one layer becomes homogeneous in *D* because isolated *C*’s (or small *C* clusters) are more easily eliminated than *D*’s. Thus, even though all four absorbing states may be reached, in principle, the probabilities to do so are very different and most likely the populations approach heterogeneous absorbing states, whereas homogeneous cooperation across lattices is a highly unlikely outcome.

## 3 Discussion

Interactions between species play a crucial role in symbiotic associations. More specifically, in mutualistic exchanges members of one species provide benefits in the form of goods or services to members of another species and vice versa. For costly provisions this results in an inter-species donation game but with a more fragile arrangement than for the standard, intra-species donation game. Here we studied the evolution of costly acts of cooperation on a two-layer square lattice, where each layer represents one species. This models a basic two-species ecological system. Evolution is controlled by increased reproduction or imitation of fitter neighbours within the same layer, while the payoff (or fitness) of individuals is determined through interactions between layers.

In recent years multi-layered networks have attracted considerable attention to model multiple, simultaneous interactions within a single species (Gómez-Gardeñes et al., 2012b; Hayashi et al., 2016; Su et al., 2022). Collective behaviour in strategy updates can also support spontaneous symmetry breaking for certain connectivity structures, such as on lattices that can be divided into two sub-lattices (Szabó et al., 2010) or two-layer networks (Gómez-Gardeñes et al., 2012a; Jin et al., 2014; Lugo et al., 2020; Takesue, 2021).

Using numerical simulations, we determined the frequency and fluctuation of cooperation in both layers as a function of the cost-to-benefit ratio, *r*. This reveals three distinct critical phase transitions: *(i)* for *r* → *r*_1_ from below, cooperation goes extinct in both layers simultaneously following the directed percolation universality class. Surprisingly, the extinction threshold *r*_1_ ≈ 0.02283(3) is very similar to the threshold in single species interactions *r* ≈ 0.0261 (estimated from (Szabó and Hauert, 2002)). Even though conditions for cooperation would seem much more challenging in interactions across species. *(ii)* spontaneous symmetry-breaking arises in the frequency of cooperation on the two layers when *r* → *r*_2_. In fact, two equivalent asymmetric phases exist where individuals in one layer prey on those in the other layer or vice versa. This phase transition is similar to the spontaneous symmetry breaking in the two-dimensional Ising model in the absence of an external magnetic field. Although, the additional degrees of freedom (four instead of two microscopic states) result in deviations in the fluctuations of cooperation frequencies. *(iii)* The third critical transition occurs for *r* → *r*_3_ from above and is driven by bursts of *DD* domains. For small *r* the magnitude of the symmetry breaking decreases with *r*, while the magnitude of bursts increases and causes fluctuations to diverge. However, the identification of the universal features of bursts and the slow transition towards absorbing states requires further, time-consuming simulations and analytical investigations (Chay et al., 1994; Vázquez et al., 2006).

From a dynamical systems perspective this last critical transition is the most intriguing feature of our model. It poses a challenge for understanding the underlying mechanisms that characterize this universality class and drive the observed power law behaviour. Conversely, from a more applied, biological or social perspective, the spontaneous symmetry breaking and the dynamics of the asymmetric phase are equally clearly the most relevant and compelling feature. In particular, the mechanism for generating and maintaining the asymmetric phase appears to be intrinsically linked to the spikes in defection observed for *r* → *r*_3_. More specifically, the emergence of asymmetries is driven by cyclical invasions of locally homogeneous domains: *CD, DC* invade *CC* invades *DD* invades *CD, DC* etc. This cycle becomes increasingly pronounced for decreasing *r* because the local domains grow and eventually give rise to the global spikes in defection that characterize the third critical transition. Assuming more *C*’s in one layer, the invading *CC* domains are more likely to encounter a *D* in the other layer, which then amplifies the asymmetry.

Symmetry breaking in the frequencies of cooperation between species has been reported before (Ezoe and Ikegawa, 2013). However, the characteristics of the species association is not merely determined by the strategy frequencies but rather by the pairings of individual strategies across layers: mutualistic (*CC* pairings), exploitative/parasitic (*CD, DC* pairings) or non-existent (*DD* pairings), i.e. no exchange of goods or services. In social dilemmas individuals fail to achieve the socially optimal outcome out of self interest. Cooperation in mutualistic interactions between species takes more varied forms than in interactions within species. Intriguingly, even in fully symmetrical mutualistic interactions, natural selection may favour asymmetric states where one species exploits the other. The resulting unfair cost allocation is reminiscent of the continuous snowdrift game in intra-species interactions, where evolution can promote the emergence of distinct cooperator and defector traits and spatial settings further promote the diversification into distinct traits (Hauert and Doebeli, 2021).

In the asymmetric phase, one species produces a social good at much higher rates than the other, which effectively separates the two species into producers and consumers. Interestingly, however, the average level of cooperation across both layers remains essentially constant at 50%. Both species produce social goods but the consumers produce theirs at much lower rates. Nevertheless, exploitation in this dynamical equilibrium is limited both spatially and temporally. In the long run, every individual produces one good and receives the other. Surprisingly, this asymmetry tends to become more pronounced for higher stakes, that is for *decreasing* cost-to-benefit ratios. Although the social dilemma becomes more benign and defection less tempting, the inequalities between species culminate in full exploitation under the most benign conditions.

## 4 Funding

C.H. acknowledges financial support by the National Science and Engineering Research Council of Canada (NSERC) Discovery grant RGPIN-2021-02608.

## 5 Data availability

The source code for the individual-based simulations and the simulation data is available at https://github.com/evoludolab/IBS-MSPD.

## Notes

### Competing Interest Statement

The authors have declared no competing interest.

